# The Functional Landscape of SARS-CoV-2 3CL Protease

**DOI:** 10.1101/2022.06.23.497404

**Authors:** Sho Iketani, Seo Jung Hong, Jenny Sheng, Farideh Bahari, Bruce Culbertson, Fereshteh Fallah Atanaki, Arjun K. Aditham, Alexander F. Kratz, Maria I. Luck, Ruxiao Tian, Stephen P. Goff, Hesam Montazeri, Yosef Sabo, David D. Ho, Alejandro Chavez

**Affiliations:** Aaron Diamond AIDS Research Center, Columbia University Vagelos College of Physicians and Surgeons, New York, NY, USA; Department of Microbiology and Immunology, Columbia University Vagelos College of Physicians and Surgeons, New York, NY, USA; Department of Pathology and Cell Biology, Columbia University Vagelos College of Physicians and Surgeons, New York, NY, USA; Integrated Program in Cellular, Molecular, and Biomedical Studies, Columbia University Irving Medical Center, New York, NY, USA; Department of Bioinformatics, Institute of Biochemistry and Biophysics, University of Tehran, Tehran, Iran; Medical Scientist Training Program, Columbia University Irving Medical Center, New York, NY, USA; Basic Sciences Division, Fred Hutchinson Cancer Research Center, Seattle, WA, USA; Division of Infectious Diseases, Department of Medicine, Columbia University Vagelos College of Physicians and Surgeons, New York, NY, USA; Department of Biochemistry and Molecular Biophysics, Columbia University Vagelos College of Physicians and Surgeons, New York, NY, USA

## Abstract

SARS-CoV-2 (severe acute respiratory syndrome coronavirus 2) as the etiologic agent of COVID-19 (coronavirus disease 2019) has drastically altered life globally. Numerous efforts have been placed on the development of therapeutics to treat SARS-CoV-2 infection. One particular target is the 3CL protease (3CL^pro^), which holds promise as it is essential to the virus and highly conserved among coronaviruses, suggesting that it may be possible to find broad inhibitors that treat not just SARS-CoV-2 but other coronavirus infections as well. While the 3CL protease has been studied by many groups for SARS-CoV-2 and other coronaviruses, our understanding of its tolerance to mutations is limited, knowledge which is particularly important as 3CL protease inhibitors become utilized clinically. Here, we develop a yeast-based deep mutational scanning approach to systematically profile the activity of all possible single mutants of the SARS-CoV-2 3CL^pro^, and validate our results both in yeast and in authentic viruses. We reveal that the 3CL^pro^ is highly malleable and is capable of tolerating mutations throughout the protein, including within the substrate binding pocket. Yet, we also identify specific residues that appear immutable for function of the protease, suggesting that these interactions may be novel targets for the design of future 3CL^pro^ inhibitors. Finally, we utilize our screening results as a basis to identify E166V as a resistance-conferring mutation against the therapeutic 3CL^pro^ inhibitor, nirmatrelvir, in clinical use. Collectively, the functional map presented herein may serve as a guide for further understanding of the biological properties of the 3CL protease and for drug development for current and future coronavirus pandemics.

## Main text

SARS-CoV-2 (severe acute respiratory syndrome coronavirus 2) has transformed daily life around the globe and caused widespread socioeconomic damage (Wu et al., 2020; Zhou et al., 2020). To curtail the ongoing pandemic, researchers have intensively studied SARS-CoV-2 in hopes of identifying effective avenues for therapeutic intervention. To that end, several antibodies and antivirals have been identified and have consequently been authorized for use in COVID-19 (coronavirus disease 2019) patients (Beigel et al., 2020; Chen et al., 2021b; Consortium et al., 2021; Gupta et al., 2021; Hammond et al., 2022; Jayk Bernal et al., 2022; Weinreich et al., 2021). Current treatments are directed towards three viral targets: the spike glycoprotein (monoclonal antibodies), the RNA-dependent RNA polymerase (remdesivir and molnupiravir), and the 3-chymotrypsin-like protease (3CL^pro^) (nirmatrelvir). Despite the promise of these therapies, previous experiences with the rapid evolution of viruses suggest that resistance is likely to quickly arise (Strasfeld and Chou, 2010). Indeed, spike variants with enhanced resistance against antibody neutralization underscores this unfortunate possibility (Iketani et al., 2022; Liu et al., 2022b; Planas et al., 2021; Wang et al., 2021a; Wang et al., 2021b).

Furthermore, studies have demonstrated that adaptation of the virus to remdesivir-mediated polymerase inhibition can be readily achieved both *in vitro* and *in vivo*, suggesting that despite the conserved nature of a viral target, resistance can still arise (Gandhi et al., 2022; Stevens et al., 2022; Szemiel et al., 2021). While no reports yet exist for SARS-CoV-2 3CL^pro^ resistance, examples of resistance to protease inhibitors in the case of HIV-1 and HCV foreshadow this likely event for the coronavirus (Clavel and Hance, 2004; Strasfeld and Chou, 2010).

The 3CL^pro^ is an attractive therapeutic target as it is essential for viral replication and there is a history of using protease inhibitors to successfully treat other viral illnesses (Chen and Njoroge, 2009; Ghosh et al., 2016; Jin et al., 2020a). To date, numerous SARS-CoV-2 3CL^pro^ inhibitors have been reported, with a few advancing into clinical trials, and one (nirmatrelvir; utilized in combination with ritonavir and marketed as PAXLOVID™ by Pfizer) being authorized for clinical use to mitigate against severe disease and deaths (Amporndanai et al., 2021; Boras et al., 2021; Dai et al., 2020; Fu et al., 2020; Hammond et al., 2022; Hattori et al., 2021; Iketani et al., 2021; Jin et al., 2020a; Jin et al., 2020b; Ma et al., 2020; Owen et al., 2021; Qiao et al., 2021; Vuong et al., 2020; Zhang et al., 2020). Such drug discovery efforts are enticing, particularly as the 3CL^pro^ exhibits significant conservation across coronaviruses, suggesting that broad-spectrum or ready-made “off-the-shelf” antivirals may be achievable for this and future epidemics. Indeed, several 3CL^pro^ inhibitors developed for other coronaviruses have been found to also hold activity against SARS-CoV-2 3CL^pro^ (Iketani et al., 2021; Resnick et al., 2021; Vuong et al., 2020). However, the tolerance of the 3CL^pro^ to mutations remains unknown; such knowledge could allow for the rational design of inhibitors with improved potency and breadth, as well as providing insight into the biology of the enzyme.

Here, we sought to systematically evaluate the plasticity of SARS-CoV-2 3CL^pro^ by interrogating the activity of all possible point mutants. We speculated that such a comprehensive analysis would reveal the malleability of the enzyme, as well as uncover conserved sites that could be used to rationally design inhibitors that are resistant to viral escape. Such extensive profiling of mutants of a gene, also known as deep mutational scanning (DMS), has been conducted successfully for the spike gene of SARS-CoV-2 by several groups, revealing the impact of mutations on expression, binding to its cognate receptor angiotensin converting enzyme 2 (ACE2), and binding by antibodies, underscoring the applicability of this approach (Garrett et al., 2021; Garrett et al., 2022; Greaney et al., 2021a; Greaney et al., 2021b; Starr et al., 2021; Starr et al., 2020).

To conduct DMS, a scalable assay that can report on the properties under study must first be designed (Fowler and Fields, 2014). We chose to work with *Saccharomyces cerevisiae* (budding yeast) due to their ease of manipulation, successful use in previous SARS-CoV-2 DMS studies, and similarity to humans with regard to their folding, post-translational modification, and turnover of proteins (Botstein and Fink, 2011; Greaney et al., 2021a; Greaney et al., 2021b; Starr et al., 2021; Starr et al., 2020). Previous reports have identified that exogenous expression of viral proteases in cells can lead to cellular toxicity (Blanco et al., 2003; Resnick et al., 2021). We speculated that a similar approach could be used with SARS-CoV-2 3CL^pro^ in yeast, and so we cloned the SARS-CoV-2 3CL^pro^ gene into an inducible yeast expression vector and generated a yeast strain containing this construct. As compared to cells containing a non-toxic control protein, enhanced yellow fluorescent protein (EYFP), cells expressing the SARS-CoV-2 3CL^pro^ showed a significant growth deficiency (**Fig. S1A**). To determine if the observed decrease in growth was due to the activity of the protease or a cryptic function performed by the exogenously expressed protease, we generated a catalytically inactive C145A mutant (Lee et al., 2020; Resnick et al., 2021). When expressed in yeast, the C145A mutant grew similarly to the non-toxic EYFP control, suggesting that the observed toxicity was a direct effect of 3CL^pro^ activity (**Fig. S1A**).

This growth defect upon expressing the 3CL^pro^ in yeast formed the basis of our assay and suggested that we could profile libraries of 3CL^pro^ mutants by growing them within a mixed pool and analyzing their relative abundances before and after inducing their expression. In this paradigm, we expected active variants to deplete and inactive variants to enrich within the pool upon induction. To generate our library of 3CL^pro^ variants, we used degenerate oligonucleotides coupled with a site-directed mutagenesis protocol (see **Methods** for details). Saturation mutagenesis of each residue was conducted by designing a single primer containing an NNK stretch, which generates all possible amino acid codings for the targeted residue. We also introduced into each primer a degenerate codon(s) upstream and/or downstream of the NNK stretch. This additional site(s) of degeneracy enabled the same amino acid mutation to be represented by a much larger number of DNA sequences than in conventional mutagenesis approaches, allowing us to average across the various DNA codings and by doing so better control for the noise inherent with assays performed at this scale (Schmierer et al., 2017; Zhu et al., 2019) (**Fig. S1B**). Once the mutant plasmids were generated, they were transformed into yeast, and cells were grown with or without induction of protease expression, followed by analysis of the abundance of the various mutants under each condition.

We conducted the DMS in biological duplicate for each of the induced and uninduced conditions. Upon observing robust correlation between the replicates, we proceeded to aggregate the effects between the two replicates, as done before (**Fig. S1C**) (Starr et al., 2020). The representation of mutations at each residue was as expected, with mostly non-synonymous mutations created by the site-directed mutagenesis method employed (**Fig. S1D**). With regard to overall coverage, we observed most variants, with 94.3% (6060 of 6426 possibilities) of all possible variants assessed for activity.

To determine the activity of each SARS-CoV-2 3CL^pro^ variant, we compared its enrichment in the induced versus uninduced conditions, and calculated an activity score (see **Methods** for details). We note that these scores by definition incorporate not just the enzymatic activity of the protease, but additional factors, such as their stability and expression. These activity scores were then normalized to the wild-type and stop coding, set at 0 and −1, respectively, and visualized as a heatmap (**Fig. 1A**). Significant heterogeneity in tolerance to mutation across the protein was observed, with some specific residues and regions appearing highly constrained and intolerant to mutation (i.e., mutations resulted in loss of activity).

**Figure 1.**
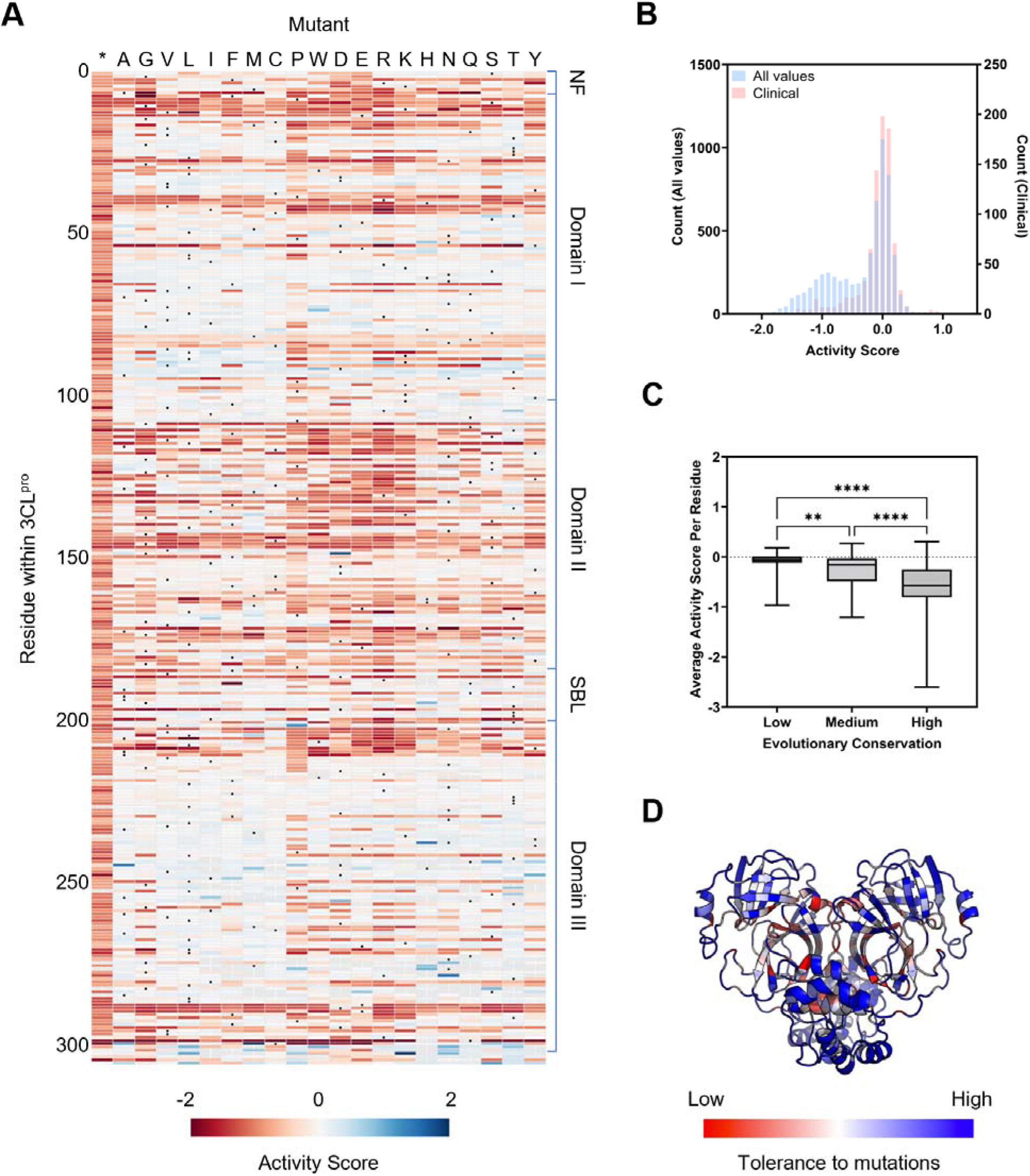
Activity of SARS-CoV-2 3CL^pro^ mutants. (**A**) Heatmap delineating protease activity of all point mutants in SARS-CoV-2 3CL^pro^. Wild-type at each residue is denoted with a black circle, asterisk designates the insertion of a stop codon at the residue. The activity score is bounded from −2 to 2. NF: N-finger; SBL: substrate binding loop. (**B**) Overlay of histograms of all activity scores in the screen and for those observed in clinical isolates. (**C**) Correlation between protease activity and evolutionary conservation of residues. Data are shown as a box and whiskers plot with whiskers denoting minimum to maximum. Dashed line is shown at 0. Statistical significance was determined by Kruskal-Wallis test followed by Dunn’s multiple comparisons test. **p < 0.01; ****p < 0.0001. (**D**) Tolerance of residues within SARS-CoV-2 3CL^pro^ for mutations overlaid onto the crystal structure.

As an initial validation of our data, we first examined known characteristics of the 3CL^pro^, either in SARS-CoV-2 or in its close relative, SARS-CoV. The 3CL^pro^ can be structurally separated into the N-finger (residues 1-7), domains I (residues 8-100) and II (residues 101-183), a substrate binding loop (SBL) connecting domains II and III (residues 184-199), and domain III (residues 200-306) (Tahir Ul Qamar et al., 2020). Looking at specific residues, mutations to the catalytic dyad, His41 and Cys145, resulted in loss of activity, as would be expected given their essential role in protease activity (Kneller et al., 2020; Lee et al., 2020). Additional critical residues within the active site were also generally conserved. These included Asp187, which stabilizes His41, and Arg40, which forms a salt bridge with Asp187 to stabilize its position (Kneller et al., 2020; Suarez and Diaz, 2020). Residues previously shown to directly bind one of the natural substrates of the 3CL^pro^, such as Phe140, Glu143, His163, His164, and Gln192, were mostly conserved and intolerant to mutations, although other substrate binding residues such as Glu166 and Gln189 were not (Lee et al., 2020; Zhao et al., 2022). These latter nonconserved residues have previously been implicated in conferring substrate flexibility for SARS-CoV 3CL^pro^, and therefore may tolerate mutation (Muramatsu et al., 2016; Zhao et al., 2022). These observations suggested that our data recapitulated known interactions and could also reveal complexities in the properties of the protease.

To further validate the data, we curated a list of mutations within the 3CL^pro^ that have been identified within patient samples and mapped them onto our DMS data (Chen et al., 2021a; Shu and McCauley, 2017). We expected that clinical variants should have high activity scores within our dataset, as viruses with inactive proteases would not be expected to replicate sufficiently to be found in clinical isolates. We identified 932 single mutants from clinical samples (with ≥3 occurrences), and found that all of these mutants clustered around the wild-type score of 0, suggesting that our data were in line with what is observed in natural infections (**Fig. 1B**).

Although the 3CL^pro^ is largely conserved, there is still variability that can be observed throughout coronavirus 3CL proteases. We speculated that residues that are less conserved across evolution should overlap with mutation-tolerant sites as denoted by our DMS data. To investigate this, we utilized the ConSurf server to determine the rate of evolution for each residue, with residues that were poorly conserved being scored as having a high rate of evolution (Glaser et al., 2003; Landau et al., 2005). We then grouped residues as having a high, medium, or low rate of evolution, and plotted the average activity score for each of the residues in the various groups. As expected, residues with high or low rates of evolution were generally tolerant or intolerant to mutation, respectively (**Fig. 1C**). Of note, while we do observe an association between these two metrics, there are additional factors that influence evolutionary conservation, such as compensatory or enabling mutations that were not considered in our screening and likely limit the strength of our association.

Finally, we considered whether our data aligned with the known structural features of SARS-CoV-2 3CL^pro^. When we mapped the average activity score at each position onto the crystal structure of the protein, we observed that surface-exposed residues were significantly more tolerant to mutations than buried residues (**Fig. 1D**). Furthermore, regions of importance, such as those forming the catalytic pocket or the dimerization interface, strongly stood out as regions that were intolerant of mutations. Collectively, the above analyses suggest that our data are robust and reflect true biological properties of the protease.

We next sought to further validate our data by investigating specific mutants indicated to be active or inactive by our screen. We chose 24 mutants throughout the gene with varying scores (14 predicted to be active and 10 predicted to be inactive), and produced isogenic yeast strains for each mutation. Each mutant was then individually tested for protease activity, and all of the mutations were found to behave as predicted from our DMS (**Fig. 2A**). Unexpectedly, various C-terminal truncations, which lack residues that are believed to be critical for protein dimerization, exhibited activity (Hsu et al., 2005). To verify this observation, we also generated an independent strain with a truncated 3CL^pro^ variant (deletion of residues 301-306). In agreement with our screening data, this truncated mutant showed protease activity (**Fig. S2A**). We note that such a mutant is likely inviable in nature in which the C-terminal residues are critical for release of the 3CL protease from the polyprotein, whereas the direct expression of the protease in our system circumvents the need for polyprotein processing.

**Figure 2.**
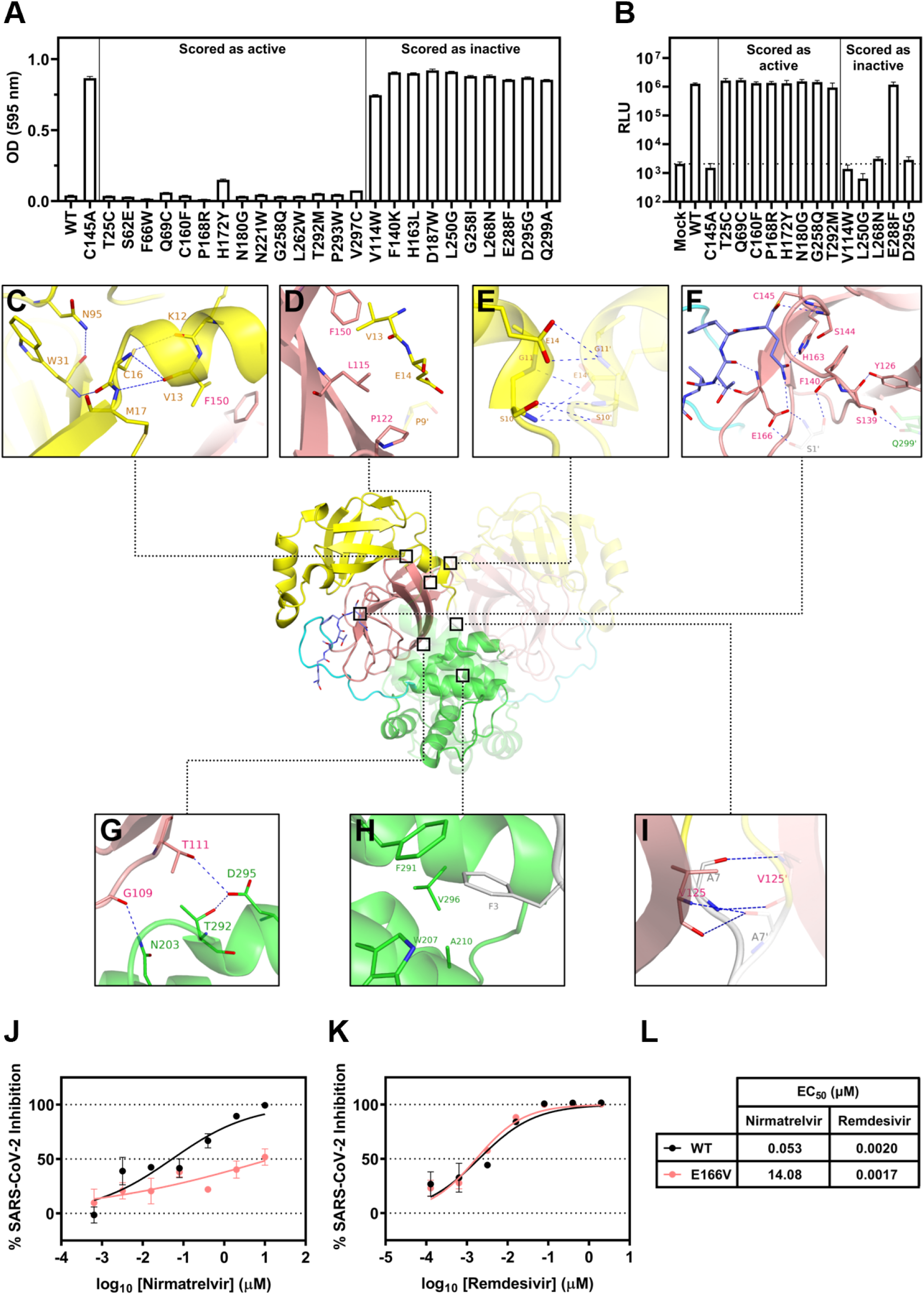
Validation of mutations and critical interactions identified in this study. (**A**) Validation in yeast of individual SARS-CoV-2 3CL^pro^ mutants that scored as active or inactive in the screen. Strains were grown for 48 h and then growth was quantified by measurement of OD at 595 nm. Data are shown as mean ± s.e.m. of technical triplicates. (**B**) Testing of recombinant live infectious virus with selected individual 3CL^pro^ mutations through a reverse genetics system. Generated mutants were used to infect Vero E6 cells for 72 h, and then infection was quantified by measurement of luciferase activity. Dashed line is shown at mean value of mock. Data are shown as mean ± s.e.m. of biological triplicates. (**C-J**) Conserved residues identified in the screen and their possible interactions and roles in 3CL^pro^ activity. The following colors are used to denote specific regions: yellow (domain I), pink (domain II), green (domain III), gray (N-finger), and cyan (substrate binding loop). The substrate is shown in dark blue. A previously solved structure of the 3CL^pro^ with its native substrate is shown (PDB: 7KHP). Residues within the other protomer are denoted with a prime symbol (‘). (**J-L**) Testing of inhibition of wild-type and E166V live infectious virus by nirmatrelvir and remdesivir. Viruses were used to infect Huh7-ACE2 cells for 24 h, and infection was quantified by measurement of luciferase activity. Data are shown as mean ± s.e.m. of technical triplicates.

While the correlation of our DMS data with clinical sequencing of SARS-CoV-2 isolates suggests that our high-throughput approach captures the true tolerance of the protease to mutation, as a further validation, we tested several mutants, as well as WT and C145A as controls, in the context of authentic SARS-CoV-2 virus. To generate our viral mutants, we used an established reverse genetics system, and introduced into it a subset of our active and inactive variants and attempted to infect Vero E6 cells (Ye et al., 2020). We found that testing within the context of the virus recapitulated the screening results, with the mutations predicted to retain an active protease resulting in a fully viable virus, whereas the mutations predicted to inactivate the protease led to inviable virus, with the exception of one mutant, E288F (**Fig. 2B**). This mutant was scored to be inactive in the screen and validated as such within yeast, yet exhibited activity within the recombinant virus. This difference in phenotype may be due to the slightly different environmental conditions between yeast and mammalian cells as well as the differences in substrates in the two contexts. Nevertheless, 14 of the 15 SARS-CoV-2 strains tested led to comparable results between the yeast and virus assays, suggesting that our screening largely recapitulates what is observed in a functional SARS-CoV-2 virus (**Fig. 2A-B**).

For the 10 inactive protease variants that were independently validated, we wondered if a loss of 3CL^pro^ dimerization, protein instability, or deficient catalytic activity might explain their lack of function. To address the question of dimerization, we designed a yeast two-hybrid assay to test whether the mutations were still competent for dimerization and found that seven of the 10 could still dimerize at varying levels, suggesting that a loss of enzymatic activity, reduction in dimerization efficiency, or a combination of both could explain the observed phenotypes (**Fig. S2B**). For the three that no longer seemed to be able to dimerize, we tested protein expression levels by immunoblotting, observing that two of the three (L250G and L268N) were poorly detected, suggesting that they formed unstable proteins. The final variant, D295G, was well-expressed, and hence its lack of signal in the yeast two-hybrid assay suggests it to be a dimerization-defective mutant, although additional experiments such as analytical ultracentrifugation are required to further investigate this mutant’s dimerization properties (**Fig. S2C**). These results suggest that despite the multiple mechanisms by which the protease can be inactivated, our DMS robustly reflects the protease activity, regardless of the nature of the mutation.

Altogether, we believe that the activity values we derived for each mutant are reflective of their true properties given the correlations we observed with clinical data, evolutionary data, and structure, in combination with the validation of individual mutations, both in yeast and authentic viruses. Therefore, we speculated that by examining residues defined to be invariant in our screen, we could identify critical interactions within the protease that could serve as targets for the next generation of protease inhibitors.

As numerous sites appeared to have varying tolerance to mutation, we focused our subsequent structural analysis on the small subset that were highly intolerant to mutations. Upon analyzing these residues within a crystal structure of SARS-CoV-2 3CL^pro^ complexed with its physiological substrate (the C-terminal autocleavage sequence, Ser301 to Gln306), we found that these sites formed critical interactions with their adjacent neighbors in three-dimensional space, both within the same or different domains of the protein, or across protease protomers (Lee et al., 2020). Within domain I, several residues likely contribute to the stabilization of the protein structure. As shown in **Fig. 2C**, the side chains of Asn95 and Cys16 form H-bonds with the backbones of Trp31 and Lys12, respectively. Backbone hydrogen bonds are also identified between Val13 and Cys16, and Met17 and Trp31. In addition, the side chains of Asn95 and Trp31 interact favorably via the π electrons of their amide and aromatic moieties, respectively. As such, Lys12, Val13, Cys16, Met17, Trp31, and Asn95 contribute to the critical intra-domain interactions to stabilize the tertiary structure within domain I, which may explain the loss of activities when these residues are mutated. Examining the interface between domain I and domain II (**Fig. 2D**), the side chains of Val13, Leu115, Phe150, Pro122, and Pro9’ of the opposing protomer form strong hydrophobic interactions. Mutating these residues to larger or more polar ones causes a reduction in protease activity, suggesting they are important inter-domain anchors. At the dimer interface between domain I and I’ (**Fig. 2E**), an extensive hydrogen bond network is observed among Ser10, Gly11, Glu14 and Ser10’, Gly11’, Glu14’ from the second protomer of the homodimer. These same residues have been shown to be critical for dimerization in SARS-CoV 3CL^pro^ (Chen et al., 2008a; Chen et al., 2008b). Looking at the catalytic domain II with a focus on the P1 pocket of the substrate binding site (**Fig. 2F**), besides the catalytic Cys145, residues including Ser144 and His163 form a multitude of hydrogen bond interactions with the substrate. The side chains of Phe140 and His163 also form a π-π stacking interaction. Moreover, Phe140 interacts with S1’ in the N-finger domain of the neighboring protomer via hydrogen bonds. This network of interactions affords the structural stability near the substrate binding site, which could explain the sensitivities of the enzymatic activities to the mutations in this region. Consistently, these residues have previously been shown to be important for SARS-CoV 3CL^pro^ catalytic activity (Barrila et al., 2006; Cheng et al., 2010; Hu et al., 2009; Huang et al., 2004; Tan et al., 2005). During close examination of our DMS data and the interface between domain II and domain III, we observed a novel interaction which our data suggest may help to stabilize this interface (**Fig. 2G**). Specifically, residues Gly109 and Thr111 of domain II form hydrogen bonds with Asn203 and Asp295 of domain III, respectively. Previous studies have implicated the importance of the N-finger in dimerization, and we observed similar results (Chen et al., 2008b; Chou et al., 2004; Hu et al., 2009; Wei et al., 2006). The N-finger folds between domains II, III, and II’ of the abutting protomer (**Fig. 2H-I**). The aromatic side chain of Phe3 is completely encapsulated in the hydrophobic pocket composed of side chains of Trp207, Ala210, Phe291, and Val296 (**Fig. 2H**). As such, the biological activity of SARS-CoV-2 3CL^pro^ is largely preserved when Phe3 is mutated to other hydrophobic residues including isoleucine, leucine, methionine, and tryptophan. However, mutations to residues of polar side chains completely abolish protein activity, corroborating the structural role of the hydrophobic encapsulation. The residue Ala7 also engages interactions at the interface of N-finger, domain II and II’ (**Fig. 2I**). Hydrogen bonds are observed between the backbones of Ala7 and Val125’. The enzymatic activities are similar to wild-type when Ala7 is mutated to valine, cysteine, serine, or threonine, but considerably lower for other mutants, suggesting the volume of the pocket is spatially constrained. In general, the structural insights support the results from our mutagenesis study, and begin to explain the mechanism by which various residues are recalcitrant to mutation.

A recent study, which explored the binding interactions of all compounds co-crystallized with 3CL^pro^ to date, identified that 185 from a total of 233 unique molecules bind within the active site, engaging the following 13 residues: Thr25, His41, Met49, Asn142, Ser144, Cys145, His163, His164, Glu166, Pro168, His172, Gln189, Ala191 (Cho et al., 2021). Our analysis (**Fig. 1A**) suggests that while some of these positions are unlikely to evolve (e.g. Ser144, His163, His172), others are more flexible (e.g. Met49, Asn142, Glu166, Pro168, Gln189, Ala191) and may therefore harbor the potential to confer resistance against many of the existing therapeutic candidates and lead compounds. This is of particular concern as the only authorized 3CL^pro^ inhibitor, nirmatrelvir (PF-07321332), is such a therapeutic, targeting the catalytic Cys145 and nestling within the substrate binding pocket (Owen et al., 2021). While most of the contact sites of this compound are intolerant to mutation, several demonstrate significant plasticity in our data. These include Glu166 at the S1 subsite, which through its side chain forms a hydrogen bond with the lactam ring of nirmatrelvir at the P1 position. From our observations, Glu166 variants that convert the acidic residue into aliphatic, basic, sulfur-containing, and even aromatic amino acids retain wild type-like activity. Residues Met49 and Gln189, which contribute to the formation of the lipophilic S2 subsite via their side chains and engage in hydrophobic interactions with the P2 dimethylcyclopropylproline group of nirmatrelvir, are also notably flexible, in accordance with other studies (Zhao et al., 2022). These analyses suggest that resistance to nirmatrelvir may readily arise.

To demonstrate this possibility, we generated a recombinant SARS-CoV-2 virus harboring the E166V mutation in 3CL^pro^ and tested its inhibition by nirmatrelvir and remdesivir as a control. As Vero E6 cells express high levels of the efflux transporter P-glycoprotein, making them unsuitable for such assays, Huh7-ACE2 cells were utilized for these assays (Owen et al., 2021). Relative to the wild-type SARS-CoV-2, inhibition of the E166V mutant by nirmatrelvir was significantly hampered, with a 265-fold loss in EC_50_ (**Fig. 2J and 2L**). In contrast, inhibition by remdesivir was unaffected, suggesting that this mutation specifically enables escape from the 3CL protease inhibitor (**Fig. 2K-L**). These data underscore the risk of resistance developing against nirmatrelvir as it becomes widely used and further highlight the value of our screen as a resource for drug design.

In summation, we have mapped the full functional landscape of the SARS-CoV-2 3CL^pro^ using a deep mutational scanning approach. Our work reveals that despite the strongly conserved nature of the gene across coronaviruses, the enzyme is highly malleable and can tolerate a plethora of mutations, even within the catalytic pocket, without significant loss of activity. These empirical results differ from bioinformatic predictions based on available sequences, emphasizing the value of this approach (Krishnamoorthy and Fakhro, 2021). Consequently, these data suggest that resistance to 3CL protease inhibitors may readily arise, and affirms the need for combinations of therapeutics to help reduce the rate of viral escape. Yet, despite this plasticity, we were able to identify highly conserved regions within the protein, both previously known and novel. Our results suggest that many of these sites are indispensable for catalytic activity or the overall structural integrity of the protein. Given their critical importance, it is natural to speculate that such residues may serve as ideal anchor points for the next generation of 3CL protease inhibitors developed to address the current and future coronavirus pandemics.

## STAR⍰Methods

### Key resources table

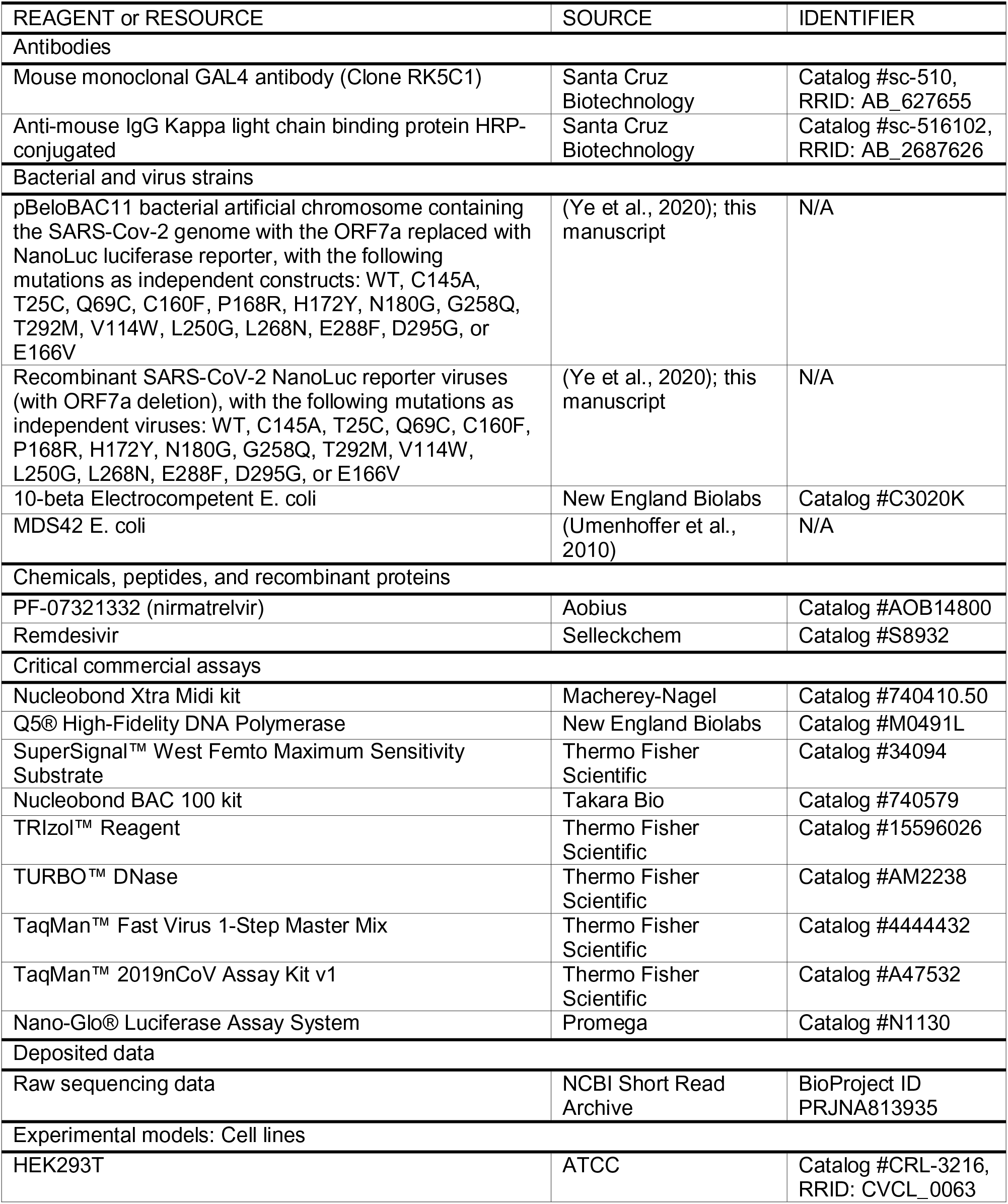
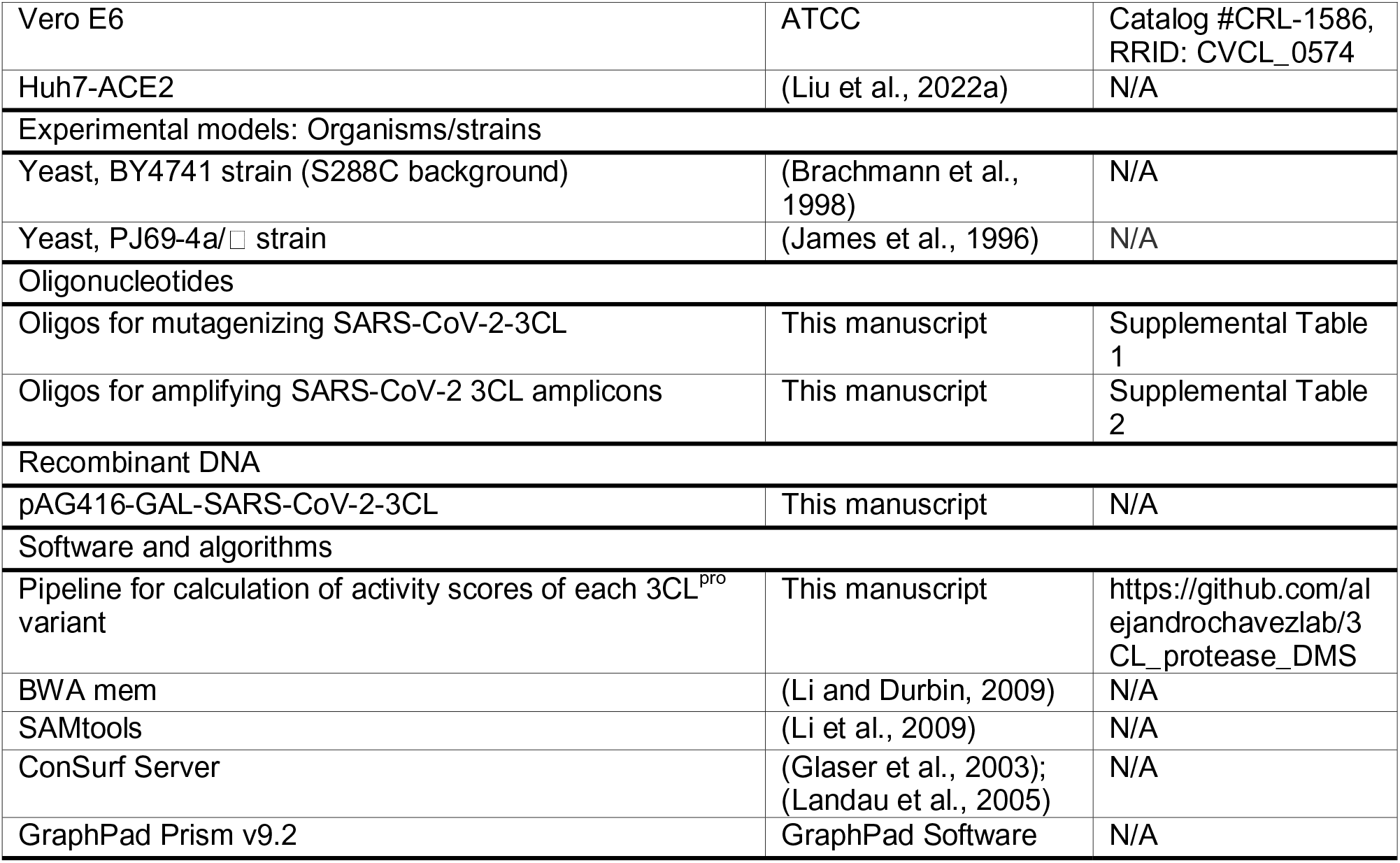

### Resource availability

#### Lead contact

Further information and requests for materials should be directed to and will be fulfilled by the lead contact, Alejandro Chavez.

#### Materials Availability

Materials used in this study will be made available under an appropriate Materials Transfer Agreement.

#### Data and Code Availability

All raw data tables and code for analysis are available at https://github.com/alejandrochavezlab/3CL_protease_DMS. The activity scores are also available as **Supplementary Table 3**. The raw sequencing data is available for download from the NCBI Short Read Archive under BioProject ID PRJNA813935.

## Acknowledgements

This work was supported by funding from the Jack Ma Foundation to Y.S., S.P.G., D.D.H., and A.C, and by funding from the JPB Foundation, Andrew and Peggy Cherng, Samuel Yin, Carol Ludwig, and David and Roger Wu to D.D.H. A.C. is also supported by a Career Awards for Medical Scientists from the Burroughs Wellcome Fund and a pilot grant from the Columbia HICCC. S.I. was supported by NIH grant T32AI106711. We thank Luis Martinez-Sobrido and Chengjin Ye for the bacterial artificial chromosome system to generate recombinant SARS-CoV-2, Jesse D. Bloom, Li Xing, and Fang-Yu Lin for help with analysis of the data, Sagi Shapira and Barry Honig for useful discussions, and Ayako Matsuda for assistance with preparation of Figure 2.

## Author contributions

A.C. conceived this project. S.I. and S.J.H. conducted the deep mutational scan. J.S., R.T., and A.C. conducted initial studies to develop the screening method. F.F., F.B., and H.M. developed and conducted the computational analyses of the screening data. J.S. and A.K.A. provided support to improve the computational analyses. S.I., S.J.H., B.C., M.I.L., Y.S., and A.C. developed the method to generate recombinant SARS-CoV-2 variants and tested those viruses. S.I., S.J.H., J.S., B.C., A.F.K., and A.C. analyzed the results. S.P.G. and D.D.H. contributed to discussions of the data and analysis. H.M., Y.S., D.D.H., and A.C. directed and supervised the project. S.I., S.J.H., B.C., and A.C. wrote the manuscript with input from all authors.

## Experimental Model and Subject Details

### Yeast

The yeast strains used for activity profiling were constructed in a BY4741 (S288C background) (Brachmann et al., 1998) and contained additional drug sensitizing mutations (Δ*pdr1* Δ*prd3* Δ*snq2*). For yeast two-hybrid studies, we used the PJ69-4a/□ background without the addition of the drug sensitizing mutations (James et al., 1996). All yeast were grown at 30 °C in a rotary shaker (for culture tubes) or in a plate shaker at 1000 RPM (for 96 deep well-plates) in growth media as appropriate.

The following yeast media were used in this study: yeast extract-peptone-dextrose medium (YPD), synthetic complete medium deficient in uracil with glucose (SC -ura GLU), synthetic complete medium deficient in uracil with galactose (SC -ura GAL), synthetic complete medium deficient in leucine and tryptophan with glucose (SC -leu - trp GLU), synthetic complete medium deficient in leucine, histidine, and tryptophan with glucose (SC -leu –his -trp GLU). YPD was formulated with 10 g/L yeast extract, 20 g/L peptone, and 20 g/L D-(+)-glucose. Synthetic complete media were formulated with 1.5 g/L yeast synthetic drop-out mix without yeast nitrogen base (US Biological), 1.7 g/L yeast nitrogen base without amino acids, carbohydrate, and without ammonium sulfate (US Biological), 5 g/L ammonium sulfate, 20 g/L D-(+)-glucose or galactose, and supplemented with the appropriate amino acids at the following concentrations: 90 mg/L histidine, 180 mg/L leucine, 90 mg/L lysine, 90 mg/L methionine, 90 mg/L adenine, and 18 mg/L uracil. For plates, the same recipe was used with the addition of 2% (w/v) agar and 600 μL of 5 M NaOH/L to help the plates solidify.

For yeast transformations, a modified lithium acetate-heat shock method was used (Gietz and Schiestl, 2007). Briefly, the appropriate parental strain was first grown to saturation in YPD, then diluted 1:100 in YPD (∼1 mL per reaction) and grown for 4 h at 30 °C to achieve log-phase growth. A plate mix was prepared by combining the following reagents per reaction: 71 µL 50% (w/v) PEG 3350, 8.8 µL 10X TE, 1.47 µL freshly boiled salmon sperm ssDNA (5 mg/mL), 8.8 µL 10X LiAc, 8.8 µL DMSO, and 1 µg of plasmid DNA to be transformed. The yeast were centrifuged at 4000 RPM for 5 min, the supernatant discarded, and then resuspended in 1X LiAc. The yeast were centrifuged again and then resuspended in 10 µL 1X LiAc per reaction. The plate mix and yeast were then combined into individual wells of a 96 well-plate. The plate was sealed and incubated for 20 min at 42 °C in a thermocycler. The plate was centrifuged at 4000 RPM for 5 min, the supernatant discarded, and then each pellet was resuspended in 20 µL sterile PBS. The yeast were then plated onto appropriate plates and grown for 48 h at 30 °C.

### Mammalian cells

HEK293T cells and Vero E6 cells were obtained from ATCC (Catalog #CRL-3216 and #CRL-1586, respectively). Huh7-ACE2 cells (Huh7 cells overexpression human ACE2) were generated previously by transduction of Huh7 cells with lentivirus encoding human ACE2, packaged using pLEX307-ACE2-blast (Addgene plasmid #158449), pMD2.G (Addgene plasmid #12259, gift of Didier Trono), and psPAX2 (Addgene plasmid #12260, gift of Didier Trono), and selecting for stable expression with 5□µg/mL blasticidin (Liu et al., 2022a). Morphology was visually confirmed prior to use and all cell lines tested mycoplasma negative.

## Method Details

### Measurement of SARS-CoV-2 3CL protease-induced toxicity

The full-length SARS-CoV-2 3CL protease gene (Genbank ID: MN908947) was synthesized (Twist Biosciences) and cloned into the yeast expression vector pAG416-GAL (Alberti et al., 2007) by Gateway cloning (Thermo Fisher Scientific). The N-terminus of the gene has the addition of a Met for translation initiation and the C-terminus is native.

Mutations were generated by site-directed mutagenesis (further detailed below). The plasmids were then transformed into the parental strain, and grown to saturation in SC -ura GLU. The yeast were then diluted 1:500 into 1 mL of SC -ura GLU or SC -ura GAL and grown at 30 °C with shaking. Growth was quantified by removing 100 µL from wells after 48 h and measuring OD at 595 nm with a spectrophotometer.

### Deep mutational scanning (DMS) of SARS-CoV-2 3CL protease

For the deep mutational scanning, libraries were prepared as biological duplicates by conducting all steps with two independent replicates.

The yeast expression vector pAG416-GAL-SARS-CoV-2-3CL was freshly prepared by Midiprep (Macherey-Nagel), and variants were introduced by a modified single primer site-directed mutagenesis method (Shenoy and Visweswariah, 2003). Oligos were designed to introduce the degenerate codon NNK at the residue of interest to introduce all possible amino acids. In addition, each oligo was designed to introduce four or more synonymous mutations at the codon(s) immediately preceding or following the residue of interest to be used as genetic redundancies to average observed effects, analogous to screening approaches in which unique molecular identifiers (UMIs) have been used (Schmierer et al., 2017; Zhu et al., 2019) (see the following **Methods** sections for further details and **Supplemental Table 1** for all oligos). Mutagenesis was conducted by mixing the following for each reaction: 150 ng template DNA, 1.25 µL 10 µM primer, 0.5 µL 10 mM dNTPs, 5 µL 5X Q5® Reaction Buffer, 0.25 µL Q5® polymerase, and H_2_O to 25 µL. All mutagenesis reactions were conducted in technical duplicate. The following cycling conditions were used:

1. 98 °C, 45 s
2. 98 °C, 15 s
3. 57 °C, 15 s
4. 72 °C, 4 min 30 s
5. Return to step #2 for 29 additional cycles
6. 72 °C, 2 min
7. Hold at 4 °C

Each reaction was then digested by adding 1 µL DpnI and incubating at 37 °C for 1 h. Technical duplicates were then combined. Multiple reactions were then further combined into sets that could be sequenced together in one amplicon (see **Supplemental Table 2** for sets). The combined reactions were then cleaned and concentrated by a PCR purification column.

Each set of mutagenesis reactions was then transformed into 10-beta electrocompetent E. coli (New England Biolabs) according to the manufacturer’s instructions. Transformations were conducted in quadruplicate to ensure adequate coverage. Following recovery and outgrowth, the bacteria was directly inoculated into liquid culture, again in quadruplicate for each transformation to avoid bottlenecking (e.g. 16 culture tubes were inoculated per set). Simultaneously, some of the bacteria was serially plated on agar plates to check transformation efficiency and to send for Sanger sequencing to confirm that mutagenesis was successful. Typical transformations resulted in 3 to 4 million colonies and 10 to 40% mutation rate when 30 colonies were sequenced.

The bacterial cultures were combined for each set and the plasmid libraries purified from these cultures by Midiprep. The libraries were transformed into the parental yeast strain. To ensure appropriate coverage, each set was independently transformed 96 times. The transformed yeast were then combined, plated on SC -ura GLU agar plates, and grown at 30 °C for 48 h. Serial dilutions were performed to check transformation efficiency. Typical transformations resulted in approximately 10,000 colonies per transformation, producing on average a million colonies across all the transformations for a given set of mutants.

Yeast were then scraped off all plates into sterile PBS. The concentration of yeast was determined by measuring OD at 595 nm with a spectrophotometer and diluted to be equivalent to a saturated culture (∼40,000 cells/µL). The yeast were then diluted 1:1000 into 1 mL of SC -ura GLU or SC -ura GAL, and grown at 30 °C for 48 h. To control against “jackpotting” events, each outgrowth condition was repeated across 24 replicates.

Following growth, DNA was extracted from the yeast using a lithium-acetate (LiOAc)-SDS lysis method (Looke et al., 2011). Briefly, plates were centrifuged at 4000 RPM for 5 min and the supernatant discarded. Pellets were resuspended in 200 µL of 200 mM LiOAc + 1% SDS, then incubated at 70 °C for 20 min. Then, 600 µL of 100% ethanol was added, and mixed well. Plates were centrifuged at 4000 RPM for 10 min, the supernatant discarded, and then allowed to air-dry until the residual ethanol evaporated. Pellets were resuspended in 200 µL of 1X TE, incubated at 42 °C for 30 min, then centrifuged again at 4000 RPM for 10 min. The supernatant, containing the extracted DNA, was removed and transferred to a new plate. DNA was stored at −20 °C until further processing.

Each set was independently amplified and indexed (see **Supplemental Table 1** for primers used for amplification of each set) and then combined and sequenced. DNA from the 24 replicates of each condition were pooled together and used as the template for first-round PCR. The following mix was used for each reaction: 0.5 µL template DNA, 0.1 µL 100 µM forward primer, 0.1 µL 100 µM reverse primer, 0.4 µL 10 mM dNTPs, 4 µL 5X Q5® Reaction Buffer, 0.1 µL Q5® polymerase, 14.8 µL H_2_O. All PCRs were conducted with six technical replicates to reduce sampling bias. The following cycling conditions were used:

1. 98 °C, 45 s
2. 98 °C, 15 s
3. 57 °C, 20 s
4. 72 °C, 1 min
5. Return to step #2 for 27 additional cycles
6. 72 °C, 3 min
7. Hold at 4 °C

PCR products were run on a gel to confirm size, and then technical replicates were combined and purified with AMPure XP beads (Beckman Coulter) according to the manufacturer’s instructions. The purified products were used as template for second-round PCR with the following mix: 0.5 µL template DNA, 0.1 µL 100 µM forward primer, 0.1 µL 100 µM reverse primer, 0.4 µL 10 mM dNTPs, 2 µL 10X Taq Reaction Buffer, 0.1 µL Taq polymerase, 16.8 µL H_2_O. All second-round PCRs were conducted in technical duplicate. The following cycling conditions were used:

1. 94 °C, 3 min
2. 94 °C, 30 s
3. 57 °C, 20 s
4. 72 °C, 30 s
5. Return to step #2 for 11 additional cycles
6. 72 °C, 3 min
7. Hold at 4 °C

PCR products were gel purified and sequenced on an Illumina NextSeq system with 75 bp single-end reads, except for Sets #8 - 10 which were sequenced on an Illumina NextSeq system with 150 bp single-end reads along with residues 113, 114, 180, 187, 196, 198, 199, and 233.

### Determination of catalytic activity of 3CL^pro^ variants from DMS

The computational pipeline for the calculation of activity scores of each 3CL^pro^ variant is available at https://github.com/alejandrochavezlab/3CL_protease_DMS. First, the raw FASTQ files were preprocessed to determine the abundances of mutations compatible with the designed oligos under each of the conditions. This was achieved by aligning the reads to the original SARS-CoV-2 3CL^pro^ sequence using BWA mem (version 0.7.17-r1188) (Li and Durbin, 2009), converting the aligned reads to bam files with SAMtools view, sort, and index (version 1.7-2) (Li et al., 2009), and then determining the counts for sequences consistent with the oligos used for mutagenesis. The raw counts for both replicates in both conditions (induced and non-induced) are available at https://github.com/alejandrochavezlab/3CL_protease_DMS.

After the preprocessing, activity scores were calculated for each variant. We first filtered out counts corresponding to sequences that differed only by one nucleotide from the native wild-type SARS-CoV-2 3CL^pro^ sequence as these counts have a possibility of being artificially inflated due to PCR or sequencing error. Additionally, we applied a count threshold of 10 for the glucose condition for each variant (no threshold for the galactose condition). Following the filtering, a pseudocount of 1 was added to all raw counts, and the resulting counts were normalized to the native wild-type read counts of each set to determine the relative abundance values of all mutant variants within the pool. Using these values, the log fold change (LFC) between the glucose and galactose conditions for each variant was calculated and an activity score determined by performing a one-sided t-test that compared the set of LFC values representing a single amino acid variant against the set of LFC values representing the recoded wild-type variant of the corresponding set. The t-statistic of the t-test was taken to be the activity score of each mutant. This t-statistic represents the difference between the LFC values of a variant and those of the wild-type variants divided by the estimated pooled standard deviation of two sets of LFC values. These obtained scores were then normalized set by set, such that wild-type = 0 and stop codon = −1. We note that the reported wild-type activity score at each residue (**Supplementary Table 3**) is derived only from the synonymous codings of the wild-type protease, and does not use the native wild-type DNA sequence which was only used for raw count normalization. The scores are available at https://github.com/alejandrochavezlab/3CL_protease_DMS and also provided as **Supplemental Table 3**.

### Identification of clinical single nucleotide 3CL^pro^ variants

To obtain a list of 3CL^pro^ variants derived from clinical specimens we made use of the COVID-CG database (Chen et al., 2021a; Shu and McCauley, 2017). The database was accessed on 3/25/2022 and used to download all amino acid variants from all geographic regions within the 3CL^pro^, corresponding to nucleotides 10,055 to 10,972. At the time the database was accessed, there were a total of 10,441,890 3CL^pro^ sequences within the database. As it is possible for interactions between mutations to convert a variant that alone is detrimental into one that is well tolerated, we curated the data provided by COVID-CG to identify mutations that occur in isolation within the 3CL^pro^ (≥ 3 occurrences), as these mutants are most similar to the variants we tested within our high-throughput study. Insertions are not considered by COVID-CG so did not need to be removed from the processed list. Deletions were removed as there is no equivalent in our data.

### Analysis of residue conservation within 3CL^pro^

Residue level evolutionary conservation was estimated by using the ConSurf server, run with the default settings on 3/25/2022 (Glaser et al., 2003; Landau et al., 2005). PDB 7JST was used as the input sequence. MAFFT was used as the MSA method. Homologs were collected from the UNIREF90 database using HMMER with settings of E-value 0.0001, 3 iterations, 95% maximal %ID between sequences, and 35% minimal %ID for homologs. 150 sequences closest to the query were chosen for sequence selection. Rate calculation was performed using the Bayesian method. Best fit was used as the model of substitution for proteins. The conservation score from 1 to 9 for each residue as calculated by ConSurf was used to group residues, with low conservation being scores 1 to 3, medium conservation being scores 4 to 6, and high conservation being scores 7 to 9.

### Yeast two-hybrid to examine protease dimerization

Yeast two-hybrid experiments were conducted by generating haploid PJ69-4a and PJ69-4□ strains expressing SARS-CoV-2 3CL^pro^ C145A with or without additional mutations and fused at the N-terminus with the Gal4 activation domain or DNA binding domain, respectively. The wild-type 3CL^pro^ could not be used for these experiments as it is toxic to the yeast when expressed (**Fig. S1A**). The MATa and MAT□ haploid strains were mated on a YPD agar plate overnight to generate MATa/MAT□ diploids, and then grown to saturation in liquid SC -leu -trp GLU media. Yeast were subsequently diluted 1:500 into 1 mL SC -leu -trp -his GLU (selection media), and grown at 30 °C. Growth was quantified by removing 100 µL from wells after 38 h and measuring OD at 595 nm to assess the dimerization capability of the mutants.

To confirm that these proteases were expressed and that the proteins were not unstable, each of the strains were also examined by immunoblotting. Strains were grown to saturation to equilibrate input cell number and then proteins were extracted from 1 mL of culture by a tricholoroacetic acid (TCA) method (Chun et al., 2019; McCullough et al., 2015). Briefly, cultures were spun down, washed once with 20% TCA, snap frozen and thawed, and then resuspended in 20% TCA with glass beads (acid-washed, 425-600 µm, Sigma). After incubation on ice for 5 min, the mixture was repeatedly vortexed and left on ice to disrupt cells, and then the pellet of extracted proteins were washed with 5% TCA. The pellet was then washed with ice-cold 100% ethanol, and then the pellet was resuspended in 1 M Tris-HCl pH 8.0. Samples were run on SDS-PAGE and transferred to PVDF membranes. Membranes were then washed, blocked with Blotting-Grade Blocker (Bio-Rad), washed, incubated overnight at 4 °C with mouse monoclonal GAL4 antibody (1:100 dilution, Clone #RK5C1, Santa Cruz Biotechnology), washed, incubated with anti-mouse IgG kappa light chain binding protein HRP-conjugated (1:1000 dilution, Santa Cruz Biotechnology) for 1 h at RT, washed, and then developed using SuperSignal™ West Femto Maximum Sensitivity Substrate (Thermo Fisher Scientific).

### Validation of 3CL^pro^ mutations in a SARS-CoV-2 reverse genetics system

Individual 3CL^pro^ mutations were introduced into the pBeloBAC11 bacterial artificial chromosome containing the SARS-CoV-2 genome in which ORF7a was replaced with a NanoLuc luciferase reporter as previously described (Ye et al., 2020). The BAC containing the wild-type 3CL^pro^ was kindly provided by Luis Martinez-Sobrido, and was propagated in the MDS42 *E. coli* strain to prevent insertion of transposable elements within the viral genome (Umenhoffer et al., 2010). To introduce 3CL^pro^ mutants, the intact BAC was digested with PacI and MluI to remove the WT 3CL^pro^ sequence, overlapping fragments were amplified using primers encoding the desired mutation (**Supplementary Table 1**), and the mutant BAC was generated by Gibson assembly (New England Biolabs). The resulting mutant BAC was electroporated into NEB 10-beta cells and purified using the Nucleobond BAC 100 kit (Takara).

These BACs were transfected into HEK293T cells using Lipofectamine 3000® (Thermo Fisher Scientific) in 12 well-plates (2 μg DNA per well) and incubated at 37 °C under 5% CO_2_ for 6-8 h. A mock transfection in which DNA was not added was included with each set of transfections. The media was changed to fresh complete media (DMEM + 10% FCS + P/S), and further incubated for an additional 40-42 h. These cells were then co-cultivated with naive Vero E6 cells in media with reduced serum (DMEM + 2% FCS + P/S) and incubated for 7 days. The supernatant was then collected, clarified by centrifugation, and aliquoted and stored at −80 °C until further use.

To normalize the amount of virus used during our subsequent assays quantifying the replication of each mutant, RNA was first extracted from an aliquot from each viral sample using TRIzol™ Reagent (Thermo Fisher Scientific) according to the manufacturer’s instructions. The extracted RNA was treated with TURBO™ DNase (Thermo Fisher Scientific), and then quantified by qRT-PCR using the TaqMan™ Fast Virus 1-Step Master Mix (Thermo Fisher Scientific) with the TaqMan™ 2019nCoV Assay Kit v1 (Thermo Fisher Scientific). These Ct values were then utilized for normalization of all of the viruses to equilibrate the input volume by RNA, such that adequate signal could be observed by the wild-type SARS-CoV-2 infection. The determined normalized volumes were used to infect naive Vero E6 cells in complete media in a 24 well-plate. After 72 h, cells were lysed and the luciferase activity was quantified using the Nano-Glo® Luciferase Assay System (Promega) according to the manufacturer’s instructions.

### Inhibition assay with recombinant SARS-CoV-2

The E166V mutant was generated as described above. Viruses were first titrated to equilibrate input. One day prior to infection, Huh7-ACE2 cells were seeded at a density of 20,000 cells per well in 96 well-plates in complete media. The following day, serial five-fold dilutions of nirmatrelvir or remdesivir were prepared in complete media, and then added to cells in triplicate. Cells were then infected with 0.05 MOI of either wild-type or E166V. After 24 h, cells were lysed and the luciferase activity was quantified using the Nano-Glo® Luciferase Assay System according to the manufacturer’s instructions. Inhibition was calculated by comparison to uninfected and infected but untreated cells. EC_50_ values were determined by nonlinear regression using GraphPad Prism v9.2.

### Statistics

P values between groups in **Fig. 1C** were determined by Kruskal-Wallis test followed by Dunn’s multiple comparisons test using GraphPad Prism v9.2.

---

**Supplemental Figure 1 – Related to Figure 1.**
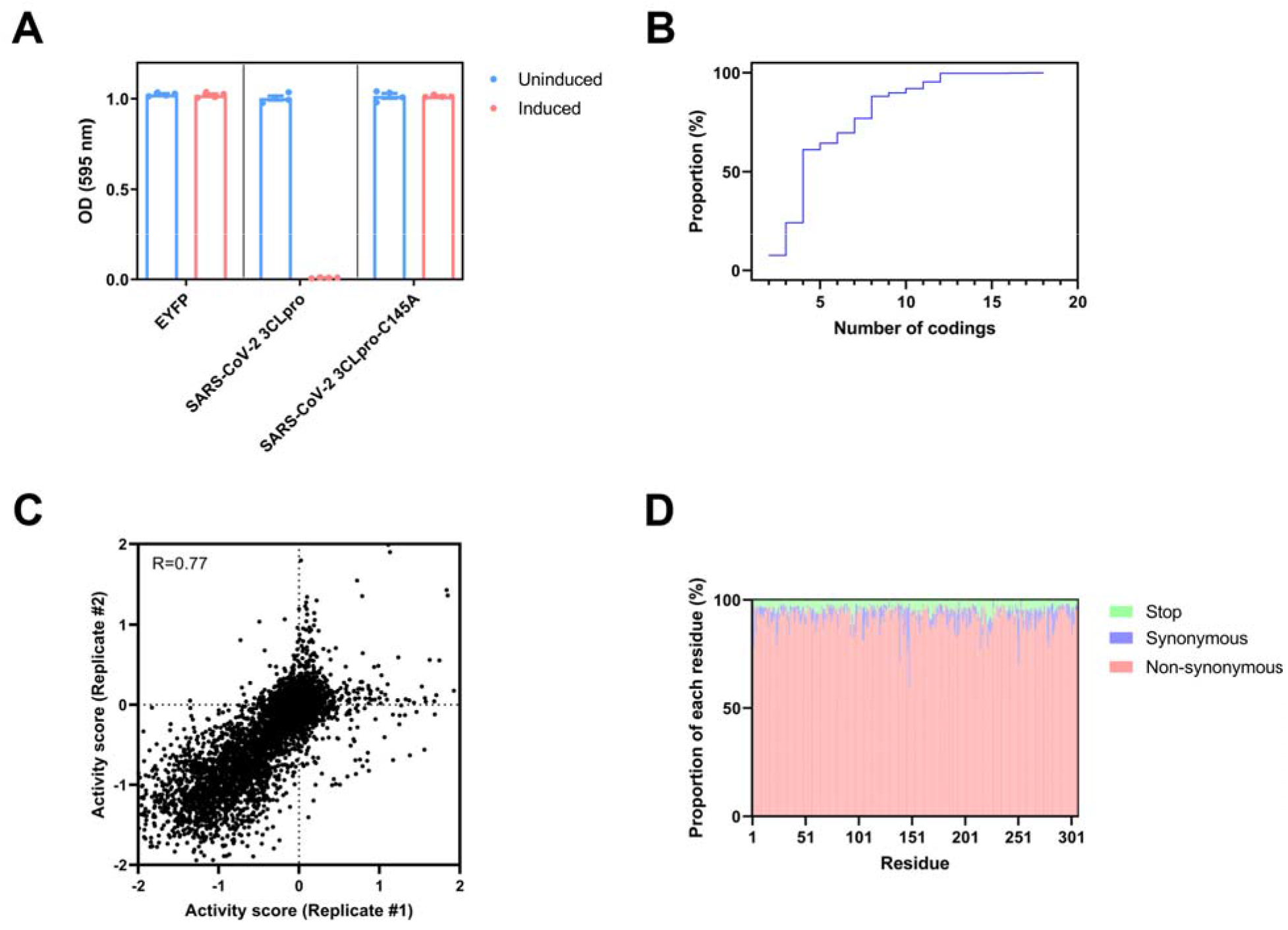
Screening system and properties of the screen. (**A**) Toxicity of SARS-CoV-2 3CL^pro^ expression in yeast. A growth defect is observed in yeast upon expression, which is dependent on the catalytic activity of the 3CL^pro^. Strains were grown in uninduced or induced conditions for 48 h and then growth was quantified by measurement of OD at 595 nm. Data are shown as mean ± s.e.m. of technical quadruplicates. (**B**) Cumulative distribution plot of codings per residue in the screen. (**C**) Correlation between biological replicates from the screen. The Pearson correlation coefficient is denoted in the top left. Axes are bounded from −2 to 2 as in Figure 1A, and outliers are therefore not shown. (**D**) Proportion of types of mutations observed at each residue in the screen.

**Supplemental Figure 2 – Related to Figure 2.**
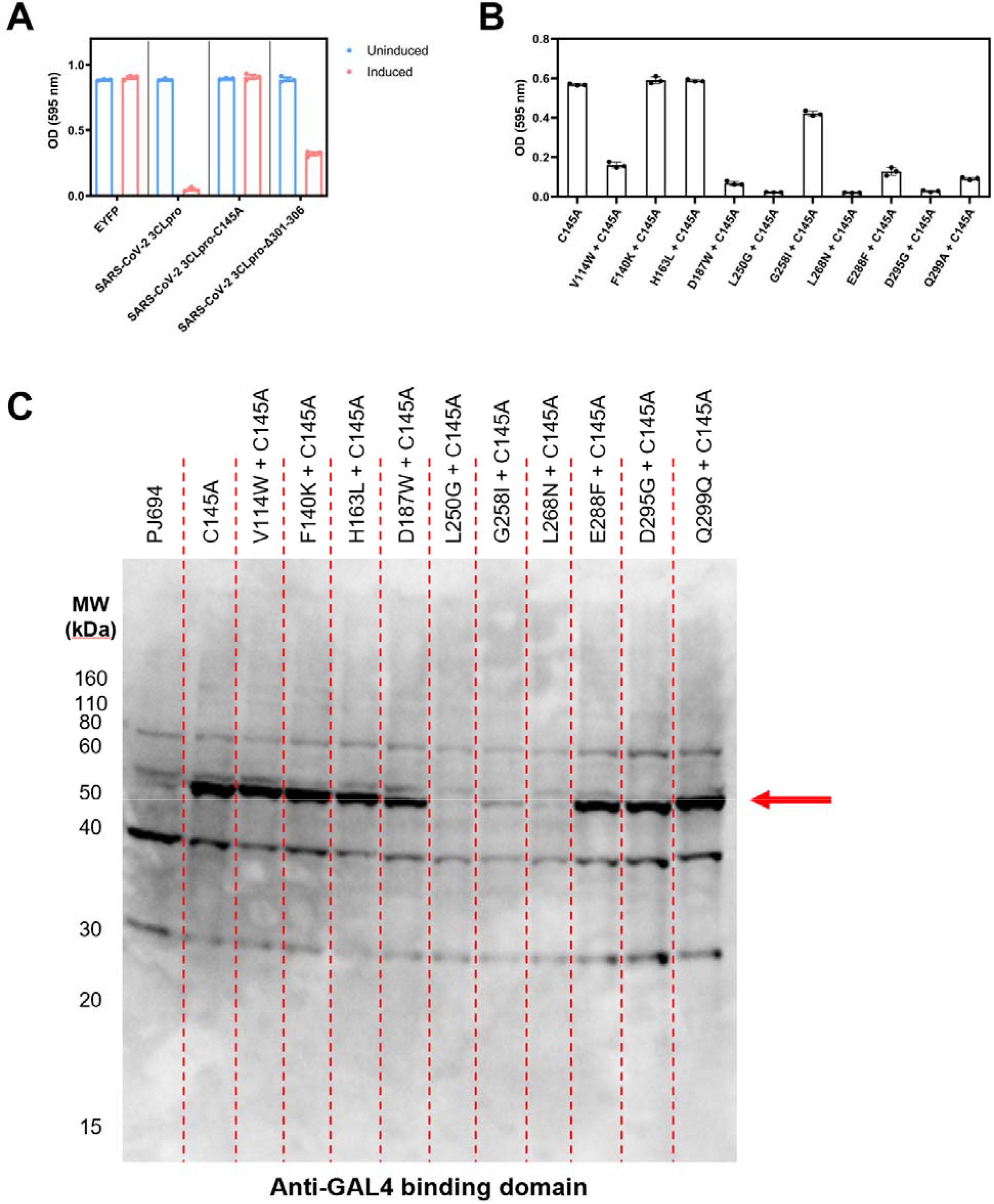
Characterization of SARS-CoV-2 3CL^pro^ mutants. (**A**) 3CL^pro^ mutant with a truncated C-terminus retains activity in this system. Strains were grown in uninduced or induced conditions for 48 h and then growth was quantified by measurement of OD at 595 nm. Data are shown as mean ± s.e.m. of technical triplicates. (**B**) Yeast two-hybrid assay to test dimerization capability of inactive 3CL^pro^ mutants. Mated strains were grown for 38 h and then growth was quantified by measurement of OD at 595 nm. Data are shown as mean ± s.e.m. of technical triplicates. (**C**) Western blot to determine if dimerization defective mutants were truly dimerization defective or if they were unable to be expressed. The band corresponding to 3CL^pro^ is indicated by the red arrow.

